# Changes in resting state functional connectivity associated with dynamic adaptation of wrist movements

**DOI:** 10.1101/2021.08.13.456278

**Authors:** Andria J. Farrens, Shahabeddin Vahdat, Fabrizio Sergi

## Abstract

Dynamic adaptation is an error-driven process of adjusting planned motor actions to changes in task dynamics (1). Adapted motor plans are consolidated into motor memories that contribute to better performance on re-exposure to the same dynamic condition. In parallel, dynamic perturbations can be compensated for by alternate motor control processes, such as co-contraction, that contribute to error reduction (2). Whether these control strategies share the same neural resources for memory formation is unclear. To address this gap in knowledge, we used a novel fMRI-compatible wrist robot, the MR-SoftWrist (3), to identify neural processes specific to dynamic adaptation and subsequent memory formation. Using the MR-SoftWrist, we acquired fMRI during a motor performance and a dynamic perturbation task to localize brain networks of interest. Resting state fMRI scans were acquired immediately before and after task performance to quantify changes in resting state functional connectivity (rsFC) within these networks. Twenty-four hours later, we assessed behavioral retention of training. A variance decomposition analysis was used to isolate behavior associated with adaptation versus alternate error reduction strategies. Immediately after the dynamic perturbation task, rsFC significantly increased within the corticothalamic-cerebellar network of the trained wrist and decreased interhemispherically within the cortical sensorimotor network. These changes were associated to behavioral measures of initial acquisition and retention, indicative of memory formation. Variance decomposition analysis revealed that increases within the cortico-thalamic-cerebellar network were associated with adaptation, while interhemispheric decreases in rsFC within the sensorimotor network were associated with alternate error reduction processes.

**Significance Statement:** Motor memory formation processes have been studied using resting state functional connectivity (rsFC) before and after exposure to adaptation tasks. However, due to technical limitations, previous studies have not investigated rsFC within brain regions localized during task execution, nor quantified rsFC within minutes after task performance. The present study used an fMRI-compatible wrist robot to localize relevant brain regions during dynamic adaptation within the cortico-thalamic-cerebellar network and bilateral sensorimotor network. Immediately following dynamic adaptation, we measured significant changes in rsFC within these networks, that were associated with adaptation behavior and with gains in behavior assessed 24 hours later, indicative of memory formation. Variance decomposition analysis identified distinct rsFC networks associated with adaptation specific processes, and with alternate motor control strategies.

---

**L**earning to control our motor actions in novel dynamic environments relies, in part, on a set of control processes that use error feedback to adjust our motor plans from movement to movement (1). Following repeated exposure to a given condition, behavior transitions from a gradual recalibration process, termed adaptation, to the automatic recall of a learned sensorimotor mapping and new behavior. The transition from adaptation to learned behavior is not well understood but is of great importance to the fields of rehabilitation, motor control and motor learning (4).

Adaptation relies on an internal model of expected task dynamics that is used to generate motor commands and predict their sensory consequences. Errors between predicted and experienced sensory feedback are used to re-calibrate the internal model and generate updated motor commands for the next movement. These adapted motor plans are consolidated into motor memories that contribute to improved performance on re-exposure to the same condition (5). In dynamic adaptation, typically studied in stable force-field environments, participants reduce their performance errors by adjusting their force profile to mirror the applied force field as they learn the new task dynamics (6). In parallel, participants can also use alternate error reduction strategies— including impedance control of their joints through muscle co-contraction— to reduce performance errors (2, 7). Currently, the neural representation of memory formation following these motor control processes is unknown.

TMS and lesion studies have identified brain regions involved in motor memory formation following dynamic perturbations immediately after task performance, that include the motor cortex (8, 9), and the cerebellum (10, 11). However, due to methodological limitations, these studies did not assess network level interactions between these regions and the rest of the brain. Functional magnetic resonance imaging (fMRI) acquired during task performance can localize brain regions associated with active motor control processes (12, 13), while resting state fMRI acquired pre- and post-task performance can measure resting state functional connectivity (rsFC) within and between these networks, reflective of motor memory formation processes (14–17). For dynamic adaptation, rsFC immediately following task performance has not been studied in networks associated with task execution due to limitations in the fMRI compatibility of robotic devices used for task performance. Moreover, while previous works have investigated contributions of perceptual learning and muscle co-contraction to changes in neural activation (16, 18), changes in neural activations specific to adaptation—independent from alternate motor control strategies—have not been determined.

This study aimed to identify the neural basis of motor memory formation processes associated with dynamic adaptation immediately following a dynamic perturbation task. We developed a novel fMRI-compatible wrist robot, the MR-SoftWrist, to study dynamic adaptation of the wrist to a lateral force field during fMRI (3). The wrist joint is optimally suited for study in the fMRI environment, as movement is distal and low-amplitude, which dually limits interference with the fMRI imaging fields and head movement during task performance. Using this device, we performed a two-day study. On day one, we localized task-related brain regions during a dynamic perturbation task, and measured rsFC immediately pre- and post task performance. On day two, we assessed participants’ behavioral retention of training.

We hypothesized that 1) rsFC within a sensorimotor and a cortico-thalamic-cerebellar network would change immediately following exposure to a dynamic perturbation condition, and that 2) rsFC measured in these networks would be predictive of behavioral measures of adaptation and of gains in performance on day two, indicative of motor memory formation. We performed a variance decomposition analysis of behavioral data to produce independent measures of adaptation and alternate error reduction strategies, that were used to identify rsFC networks specifically associated with adaptation or alternate error reduction strategies.

## Results

Data were collected from 30 participants (19 M, 11 F, age: 23 ± 4 years), who completed two experimental sessions, conducted 24 hours apart. The experimental protocol is shown in Fig. 1. The first session, conducted on day one, comprised a motor performance task (zero force condition) performed using the MR-SoftWrist to assess behavior and neural activations associated with task execution in the absence of motor learning. This served as a control condition for changes observed in a dynamic perturbation task (lateral force condition), in which the MR-SoftWrist applied velocity-dependent lateral forces to enable investigation of behavior and neural activity associated with dynamic motor control. Throughout all conditions, error clamp trials (force tunnel condition) were applied unexpectedly to enable measurement of participants’ prediction of required task dynamics (19). The MR-SoftWrist recorded participant kinetics and kinematics during task performance, while fMRI was used to measure neural activity associated with task performance. Resting state scans were acquired before and after each motor task, and behavioral retention of training was assessed 24 hours later in a repeated dynamic perturbation task.

**Fig. 1.**
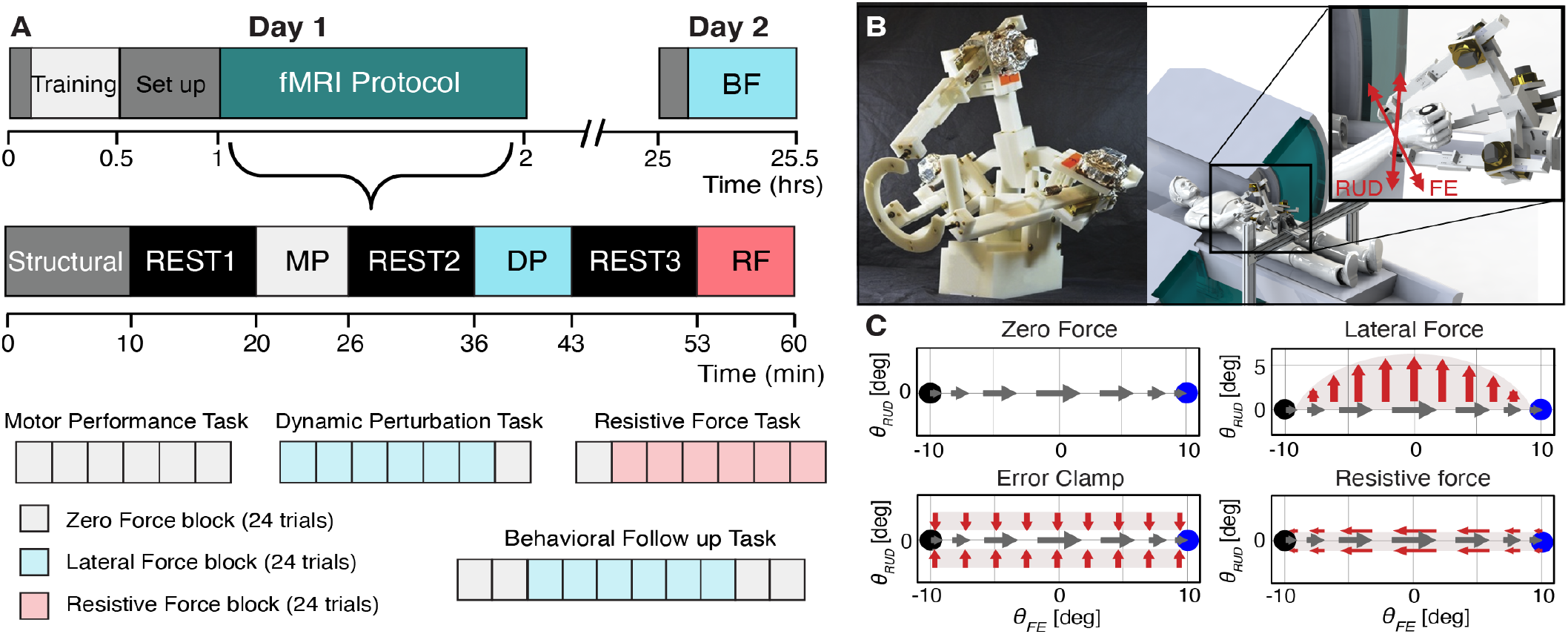
A: Experimental protocol. On day 1, participants performed three motor task with the MR-SoftWrist in the scanner, interleaved with resting state functional scans. Motor tasks included a motor performance task (MP), executed in a zero force environment, a dynamic perturbation task (DP), executed in a velocity-dependent lateral force field environment, and a resistive force task (RF), executed in a velocity-dependent resistive force environment. Prior to the fMRI protocol, participants were trained on interaction with the robot and task instructions using the motor performance task. Resting state scans were acquired to assess baseline functional connectivity (REST1), functional connectivity following motor performance (REST2) and functional connectivity following motor adaptation (REST3). On day 2, a behavioral follow-up task (BF) was performed to assess change in behavior in the lateral force field environment that would indicate retention of day one training (i.e. motor memories were formed). B: The MR-SoftWrist and a CAD rendering of the device in its operating condition in the scanner. Red arrows depict the rotational axes supported by the device that correspond to flexion-extension (FE) and radial ulnar deviation (RUD) of the wrist. C: Task-specific force fields used in the experimental protocol. Grey arrows depict exemplary cursor velocity during task execution, while red arrows depict the corresponding direction of robot applied forces.

### Behavior

Individuals can use at least two motor control strategies to reduce performance errors caused by a lateral force field, that correspond to different ways of adjusting their motor plan:

1. through adaptation of the planned force profile executed during their movements (i.e., applying forces in the opposite direction of the applied force field, which crucially requires creating an internal model of the new dynamic condition) or
2. through the use of alternate error reduction strategies, including co-contraction of the wrist muscles to modulate joint impedance to reject force perturbations (2). These control processes can occur in parallel. In our dynamic perturbation task both strategies result in a reduction of performance error. To quantify performance error in all conditions, we calculated the angular trajectory error on each trial. To quantify wrist kinetics, we calculated participants’ adaptation index during error clamp trials (interspersed throughout all task conditions with a 1/8 frequency). Across the dynamic perturbation task, reductions in trajectory errors reflect both adaptation and alternate motor control strategies. Instead, adaptation-specific recalibrations of the internal model are measurable as “after-effects” in trajectory errors that occur in the opposite direction of the applied force field when the force field is removed, and as increases in adaptation index across the lateral force condition that persist temporarily when the force field is removed.

We used two repeated measures ANOVAs to test for an effect of experimental phase on trajectory errors and adaptation index. For trajectory error, we investigated the effect of ten experimental phases including Baseline, Initial lateral force errors (trials 2-4), Mid lateral force errors (trials 4-24), Late lateral force (LF) errors, and After Effects on both Day 1 and Day 2 (Fig. 2). The ANOVA identified a significant effect of experimental phase on trajectory errors (DF: 9, F Ratio: 397.36, *p <* 0.001, *R*_*adj*_^2^ = 0.930). Post-hoc Tukey testing showed a significant increase in Initial Errors over Baseline (*p <* 0.001), that decreased across the lateral force condition on both days (Mid Errors < Initial Errors, *p <* 0.001; Late LF < Mid Errors, *p <* 0.001), consistent with improvements in task performance due to adjustments in dynamic motor control. On both days, there were significant After Effects compared to Baseline (*p <* 0.001), indicative of adaptation. Between days, contrasts of parameter estimates showed there were significantly smaller initial errors (IE) on Day 2 compared to Day 1, consistent with “recall” (Day 2 IE − Day 1 IE: − 2.379 ± 0.92 deg, *p* = 0.0103, Fig. 2) (20), and significantly smaller final errors on Day 2 compared to Day 1 (Day 2 Late LF − Day 1 Late LF: − 1.862 ± 0.92 deg, *p* = 0.0441). There was no significant difference in mid-errors between days, that would be indicative of effects in relearning, or in the magnitude of after-effects. For adaptation index, we defined eight experimental phases, including Baseline, Early lateral force, Late lateral force, and After Effects on both Day 1 and Day 2 (Fig. 2). The ANOVA identified a significant effect of experimental phase (DF: 7, F Ratio: 49.54, *p <* 0.001, *R*_*adj*_^2^ = 0.639). Post-hoc Tukey testing showed a significant increase in all experimental phases compared to Baseline (*p <* 0.001), that increased over the course of the lateral force condition on both days (Late LF > Early LF, *p <* 0.001), consistent with adaptation (Fig. 2, Bottom left). Contrasts performed between parameter estimates of experimental phases on Day 1 and Day 2 returned a greater adaptation index in Late LF on Day 2 compared to Day 1 (Day 2 Late LF − Day 1 Late LF: 0.0575 ± 0.0299 n.u., *p* = 0.0558), in line with some retention of Day 1 training. There were no significant differences between days in Early LF, or after effects.

**Fig. 2.**
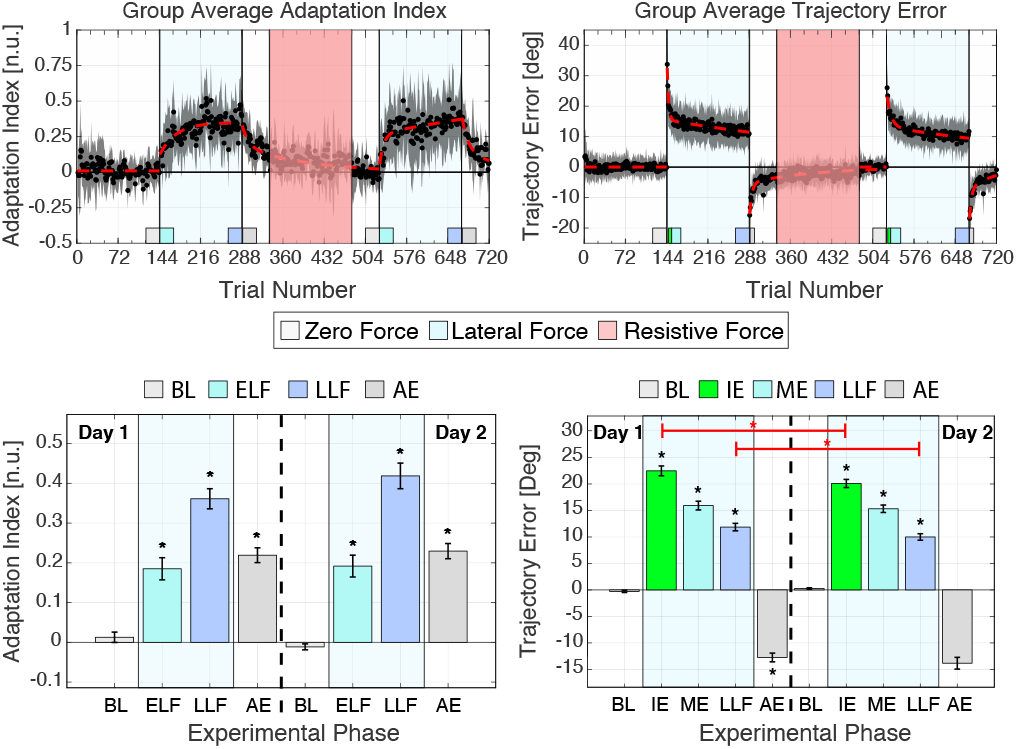
Top left: Group average adaptation index. Black dots denote the average adaptation index measured on each trial. Grey shaded regions represent the s.e.m.. Top Right: Group average trajectory error. Bottom left: Bar plot of mean adaptation index measured in each experimental phase of interest. For space, experimental phases have been abbreviated as follows: Baseline as BL, Early LF as ELF, Late LF as LLF, and After Effects as AE. Error bars depict standard error across subjects within each phase. Black asterisks indicate phases that are significantly different from baseline behavior, while red asterisks denote significant changes in measured behavior between days (*p <* 0.05). Bottom Right: Bar plot of the average trajectory error measured in each experimental phase of interest. For space, experimental phases have been abbreviated as follows: Baseline as BL, Initial Errors as IE, Middle Errors as ME, Late LF as LLF, and After Effects as AE.

#### Subject-specific Behavioral Metrics

To determine associations between behavior and rsFC, we created subject-specific behavioral measures of participants’ relative error reduction and magnitude of adaptation by fitting a double exponential model to individual subject’s trajectory error and adaptation index data. Relative error reduction (RER) was defined as the difference between model estimated final and initial trajectory errors, divided by the initial error, and had an average model fit to data in the lateral force condition of *R*^2^ = 0.300 ± 0.0055 (mean ± s.e.m.) for Day 1, and *R*^2^ = 0.303 ± 0.0048 on Day 2. Magnitude of adaptation index (MAI), defined as the model estimated end magnitude of adaptation index in the lateral force condition on each day, had an average fit of *R*^2^ = 0.386 ± 0.0058 on Day 1, and *R*^2^ = 0.397 ± 0.0065 on Day 2. For estimation of magnitude of after-effects (MAE), defined as the additive inverse of the model estimated initial trajectory error in the zero force condition following the lateral force condition, the average model fit across subjects was *R*^2^ = 0.464 ± 0.0068 on Day 1, and *R*^2^ = 0.463 ± 0.0071 on Day 2. The additive inverse was used to match the sign of the MAI metric.

We expected MAE and MAI, which both quantify adaptation specific processes, to be highly correlated. These metrics were correlated with an *R*^2^ = 0.25, *p <* 0.005 for Day 1, and with an *R*^2^ = 0.53, *p <* 0.001 for Day 2. If error reduction is achieved by adaptation, we expect metrics of adaptation magnitude to explain measures of relative error reduction. On day one, RER was significantly correlated with MAI (*R*^2^ = 0.28, *p <* 0.003) and MAE (*R*^2^ = 0.13, *p <* 0.05). For day two, RER had a positive, but not significant, association to MAI (*R*^2^ = 0.11, *p* = 0.076) and MAE (*R*^2^ = 0.11, *p* = 0.075). Such a weak association could be influenced by retention of day 1 learning, that may result in lower relative error reductions paired with greater adaptation due to recall and storage of the internal model from prior training. Or it may be due to individuals using a different mix of control strategies in response to dynamic perturbations, as RER quantifies both aspects of the motor control response while MAI and MAE predominantly capture effects of adaptation.

#### Variance Decomposition Analysis

To isolate behavioral effects associated specifically with adaptation from those associated with alternate error reduction strategies, we performed a two-part variance decomposition analysis using our behavioral summary metrics (RER, MAE, and MAI), to create a metric of adaptation specific control processes for each subject (MAD), and an independent metric quantifying alternate error reduction strategies (AERS). For day 1 behavior, AERS was strongly correlated with RER (*R*^2^ = 0.718), and mostly orthogonal to MAI (*R*^2^ = 0.003) and MAE (*R*^2^ = 0.006). Instead, MAD was significantly correlated with all three measures (RER *R*^2^ = 0.282; MAI *R*^2^ = 0.841; MAE *R*^2^ = 0.647), as expected. For day 2, AERS was again correlated with RER (*R*^2^=0.718), and orthogonal to MAI (*R*^2^ = 0.002) and MAE (*R*^2^ = 0.0001). For day 2, MAD was significantly correlated with both adaptation metrics (MAI *R*^2^ = 0.683; MAE *R*^2^ = 0.977), but not significantly associated with RER (*R*^2^ = 0.119, *p* = 0.062).

### Changes in rsFC

We aimed to determine if resting state functional connectivity (rsFC) was modulated by motor task performance within brain networks engaged during active task performance. We hypothesized that there would be no significant change in rsFC following our motor performance task, as limited learning occurs within this task, and that there would be significant change in rsFC following the dynamic perturbation task due to motor adaptation, and subsequent motor memory formation.

We used task-based fMRI scans to determine relevant brain regions of interest (ROIs) for use in an ROI-based analysis of our resting state data. This resulted in a total of 65 ROIs, including the bilateral primary motor cortices (M1), premotor cortices (PMv, PMd, SMA), primary sensory cortices (S1, BA1-3), superior and inferior parietal lobules (SPL5, SPL7, Anterior IPL, Posterior IPL), secondary somatosensory cortices (SII), intraparietal sulci (IPS), angular gyri, Broca’s Areas 44 and 45, the anterior cingulate and basal ganglia (putamen, caudate), thalamus, insular cortices, frontal pole regions (BA8, 9, 10, 46), cerebellar lobules IV-VII, crus I and II, and the anterior and posterior vermis. For the left cortical areas in the sensorimotor network, which are highly engaged during task execution, we defined subject-specific ROIs based on task related activation measured within each anatomical region of interest. For ROIs included in our restricted network analysis (detailed below), we created a single ROI within the left primary motor cortex, premotor cortex and primary sensory cortex (L-M1 (mean ± s.e.m. [range]): 184.5 ± 17.2 [136, 222] voxels; L-PM: 176.0 ± 13.0, [155, 212] voxels; L-S1: 188.4 ± 15.7 [160, 212] voxels). For our expanded network analysis, L-M1 was defined the same, however subject specific ROIs were created in each sub region within the premotor (PMv, PMd, SMA) and primary sensory cortex (BA1, BA2, BA3): (L-PMv: 193.6 ± 9.5, [167, 208] voxels; L-PMd: 181.1 ± 15.5 [160, 212] voxels; L-SMA: 174.7 ± 28.4 [63, 212] voxels; L-BA1: 167.7 ± 27.4 [99, 195] voxels; L-BA2: 167.3 ± 31.7 [70, 196] voxels; and L-BA3: 191.3 ± 16.0 [146, 236] voxels). For all other ROIs included in our analyses, we used their anatomical definition.

#### Restricted Network Analysis

From the larger network of 65 task related ROIs, we selected a sub-set of ROIs based on prior knowledge of brain networks associated with adaptation (16, 20, 21) to use in a restricted ROI-based analysis. These ROIs included the bilateral primary motor cortices (M1), premotor cortices (PM), and primary sensory cortices (S1), the left thalamus, and the right cerebellar lobules 6 and 8 (CB6, CB8). The six bilateral cortical ROIs (L-M1, L-S1, L-PM, R-M1,R-S1, R-PM) formed a cortical sensorimotor network of interest while the left motor areas, left thalamus, and right cerebellar ROIs formed a restricted cortico-thalamic-cerebellar (CTC) network of the trained wrist (L-M1, L-PM, L-S1, L-Thalamus, R-CB6, R-CB8). Both networks were formed as a complete graph with 6 nodes and 15 edges. RsFC between nodes was defined as the Fisher-transformed correlation coefficient of the average BOLD signal measured within each ROI. Within each network, we performed a mixed model analysis to test for the effect of condition (REST1, REST2 and REST3) on measured rsFC for each edge in each network of interest.

The mixed model identified significant changes in rsFC following both the motor performance and dynamic perturbation tasks (Fig. 3, Left). Following motor performance, the model identified decreases in rsFC between nodes in the left and right hemisphere of the sensorimotor network, and an increase in rsFC between the left primary motor cortex and the left thalamus (Model *p*_*FWE*_ *<* 0.05: L-PM to R-M1: *p* = 0.014, L-M1 to L-Thalamus, *p* = 0.0032; Model *p*_*unc*_ < 0.01: L-PM to R-S1: *p* = 0.005). Following the dynamic perturbation task, there were increases in rsFC within the trained cortico-thalamic-cerebellar network, including increased rsFC between left cortical regions to both the thalamus and cerebellum (Model *p*_*FWE*_ < 0.05: L-M1 to L-thalamus: *p* = 0.0016, L-S1 to R-CB6: *p* = 0.0006; Model *p*_*unc*_ < 0.01: L-M1 to R-CB6: *p* = 0.001), as well as continued decreases in connectivity between nodes in the right and left hemispheres of the sensorimotor network (Model *p*_*FWE*_ *<* 0.05: L-PM to R-M1: *p* = 0.0009; Model *p*_*unc*_ < 0.01: L-PM to R-S1: *p* = 0.002, L-M1 to R-M1: *p* = 0.001). Contrasts between REST3 rsFC and the average of rsFC measured in REST1 and REST2, executed to control for changes in functional connectivity following the motor performance task, returned significant changes in rsFC in the same edges as the contrast between REST3 and REST1, except for left primary motor cortex to the left thalamus (Model *p*_*FWE*_ < 0.05: L-S1 to R-CB6: *p* = 0.0032, L-PM to R-M1: *p* = 0.0138; Model *p*_*unc*_ < 0.01: L-M1 to R-M1, *p* = 0.007, L-M1 to R-CB6, p =0.002).

**Fig. 3.**
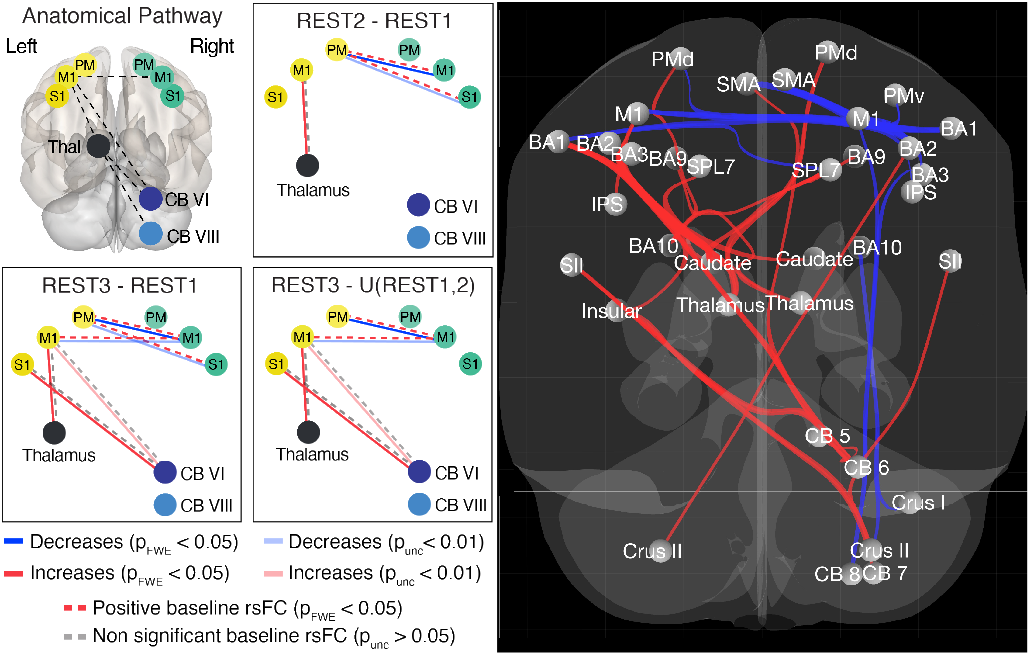
Left: Restricted Network Analysis results. Decreases in rsFC between ROIs are depicted in blue, while increases in rsFC are depicted in red. Model results with *p*_*FWE*_ < 0.05 are reported in solid (100% opaque) lines, while results with *p*_*unc*_ < 0.01 are depicted by faded (50% opaque) lines. Right: Expanded Network Analysis results. Significant network level results from network-based statistical analysis of REST3*−*REST1, projected on a glass brain (t-voxel > 3, p-cluster-FDR < 0.05). Analysis returned two networks, a sensorimotor network and a cortico-thalamic-cerebellar network, that comprised a total of 35 ROIs and 84 edges, from our original network of 65 ROIs and 4160 edges.

#### Expanded Network Analysis

To supplement our restricted ROI-based analysis, we performed a network-based statistical analysis (22) to determine if rsFC significantly changed between resting state conditions in any sub-networks of ROIs within our entire task-related ROI network (65 ROIs). Our network-based statistical analysis showed no significant changes following the motor performance task (REST2 − REST1), in line with our hypothesis. A contrast between the REST3 and REST1 conditions supported results found in our restricted network analyses, with significant (*T >* 3, *p*_*FDR*_ < 0.05) decreases in interhemispheric rsFC within a bilateral sensorimotor network, and significant increases in rsFC within the cortico-thalamic-cerebellar network (Fig. 3, Right). These results remained significant for a contrast between connectivity scores measured in REST3 and the average of scores measured in sessions REST2 and REST1.

#### ROI-to-Voxel Analysis

Finally, to supplement our ROI-ROI analyses, we performed a GLM analysis of our three ROIs in the left sensorimotor network (L-M1, L-S1, L-PM) to resting state activity measured across the whole brain in each rest condition. Group level contrasts were used to identify changes in rsFC between conditions. Our analysis identified significant positive baseline (REST 1) rsFC between the left sensorimotor ROIs (L-M1, L-PM, L-S1) and bilateral motor areas, the frontal cortex, the basal ganglia and the cerebellum (regions V-VI, VIII). For the REST2 − REST1 contrast, no significant changes were returned for the left M1 or left PM ROIs. The left S1 ROI had significant increases in rsFC to the visual cortex. For the REST3 − REST1 contrast, the left PM ROI showed significant decreases to the right primary and sensory motor cortex (BA4, BA3, BA1). The left M1 ROI showed significant decreases to the right primary motor and sensory motor cortex (BA4, BA1, BA3), increases to the left thalamus and left caudate, and increases to the right cerebellar lobules 6 and 8 (Fig. 4). The left S1 ROI showed significant decreases to the superior parietal lobe (SPL7, IPL), and significant increases to the left thalamus and caudate, and the right cerebellar lobule 6. Both the left M1 and S1 ROIs showed significant increases to the visual cortex.

**Fig. 4.**
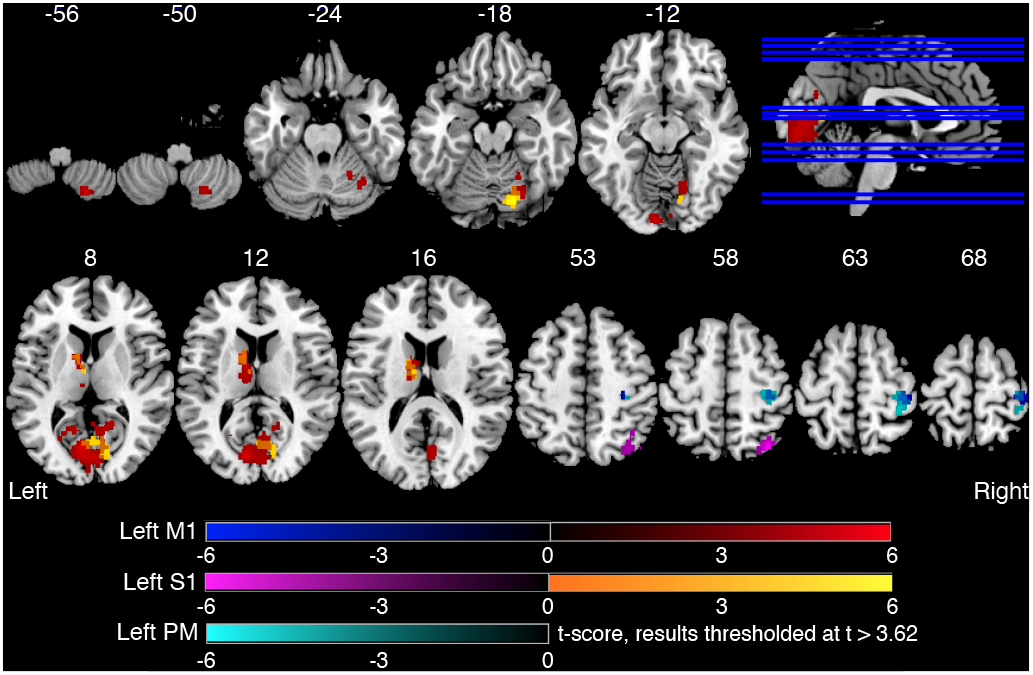
ROI-to-Voxel Analysis results. Change in rsFC between each left sensorimotor seed (M1, PM, S1) to whole brain activity following the dynamic perturbation task (REST3*−*REST1). Each statistical parametric map is overlaid on axial slices of the standard MNI 152 template, with z coordinates reported in mm. For each ROI, the seed-voxel T-maps were thresholded at *p <* 0.0011, |*T* (29)| > 3.62 at the voxel level, with FDR correction applied at the cluster level (*p*_*FDR*_ *<*0.05, K *≥* 90)

### Associations between rsFC and behavior

To investigate associations between behavior measured on day 1 and rsFC measured in each resting state condition, we performed a linear regression between behavior (MAE, MAI, RER) and rsFC measured within our two restricted networks of interest (sensorimotor and cortico-thalamic-cerebellar). For associations to day 2 behavior, we performed the same linear regression analysis, but included a term for day 1 behavior to account for effects described by initial task performance. For edges with rsFC that was significantly associated to behavior in our linear regression analysis, we performed an exploratory analysis using the metrics from our variance decomposition analysis to determine if observed effects in our rsFC conditions (REST1, REST2 − REST1, or REST3 − REST1) were driven by adaptation (MAD), or alternate error reduction strategies (AERS).

Within the sensorimotor network, 3 out of 15 edges had significant associations to behavior, while 5 out of 15 edges had significant associations to behavior in the cortico-thalamic-cerebellar network (Fig. 5). Within the sensorimotor network, all 3 edges were associated with day 1 behavior (2 to RER, 1 to MAE), and none were associated with changes in day 2 behavior. These effects were associated with alternate error reduction strategies (Fig. 5, right). Within the cortico-thalamic-cerebellar network, one edge showed a significant association with day 1 adaptation (MAE), while all 5 nodes showed significant associations to day 2 behavior when controlling for day 1 behavior (2 to RER, 2 to MAE, 3 to MAI, with overlap). These effects were preferentially associated with adaptation specific processes (Fig. 5, right). A network level breakdown of these effects by resting state condition is provided below.

**Fig. 5.**
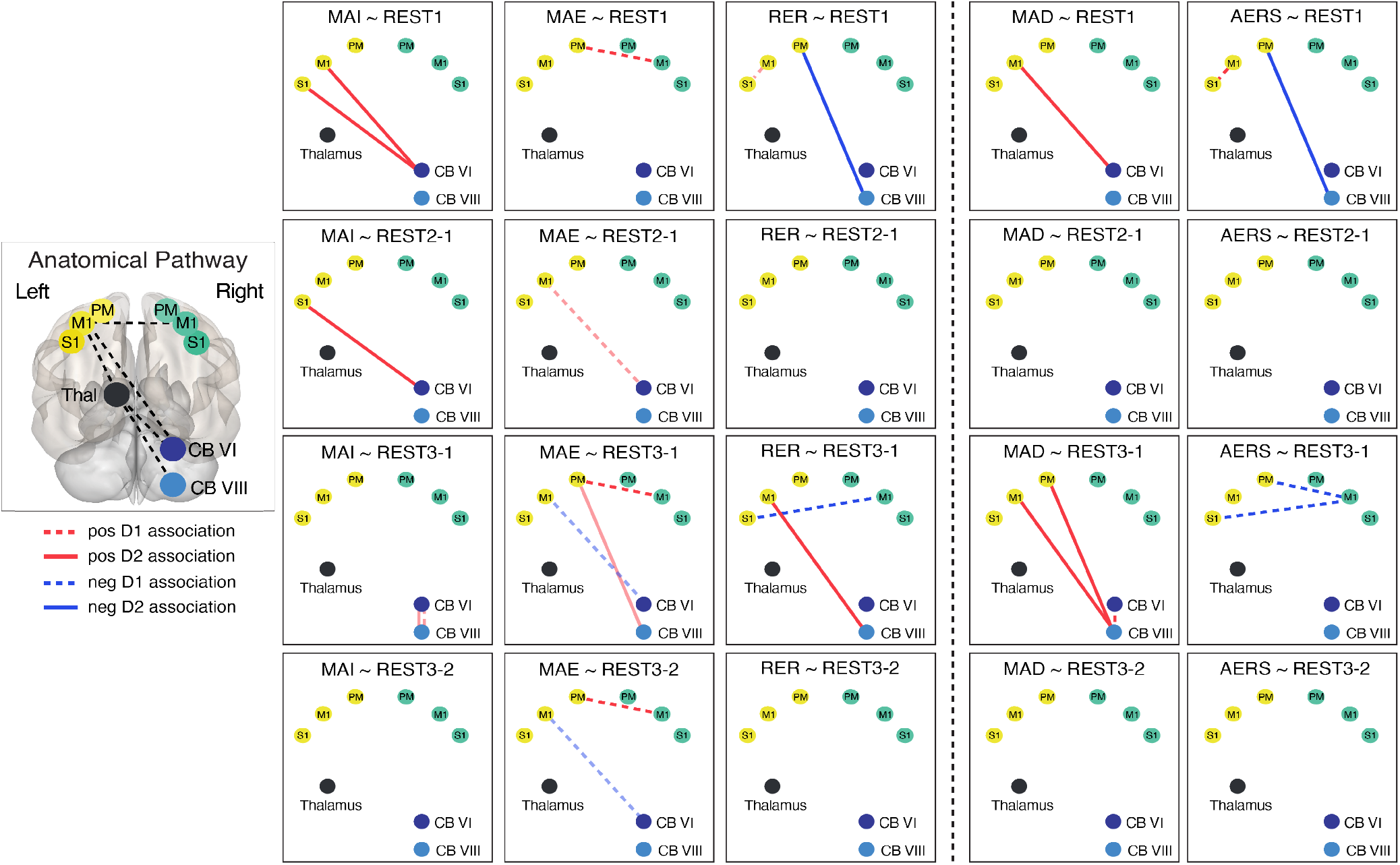
Left: Association between rsFC and D1 behavior (dashed lines) and between rsFC and Day 2 behavior (solid lines). Edges with significant association (*p*_*FWE*_ < 0.05) as determined in the reduced model analysis are depicted as opaque lines, while edges with associations at *p*_*unc*_ < 0.01 are shown as faded lines with 50% opacity. For both thresholds, only terms with *p*_*β,corr*_ < 0.05 are shown. Right: Variance decomposition analysis results. Edges with a significant association (*p <* 0.05) to AERS or MAD are reported. For space, REST3*−*REST1 has been abbreviated as REST3-1, REST2*−*REST1 as REST2-1, and REST3*−*REST2 as REST3-2

#### Associations between behavior and rsFC in the sensori-motor network

Within the sensorimotor network, linear regression analysis returned no significant (*p*_*FWE*_ < 0.05) associations between baseline rsFC (REST 1) and day 1 behavior. At the *p*_*unc*_ < 0.01 level (not corrected for family wise error) a positive association between relative error reduction and baseline rsFC was identified within the trained left sensorimotor cortex (RER ∼ L-M1 to L-S1, *p*_*model*_ = 0.009, *p*_*REST* 1_ = 0.009, Tbl. 1). The variance decomposition analysis returned an association between rsFC in this edge and alternate error reduction strategies (*p* = 0.0097). There were no significant associations between baseline rsFC and day 2 behavior, nor between change in rsFC across the motor performance task (REST2 − REST1) and behavior on either day.

**Table 1.** Top: Results of model based analysis of rsFC to behavior (RER, MAE, MAI) measured on day 1 and day 2. The table reports the most reduce model parameter estimates and their significance. Model significance is bolded for *p*_*FWE*_ *<*0.05, and non-bolded for *p*_*unc*_ < 0.01. Individual model terms are bolded for *p*_*β,corr*_ < 0.05. Bottom: Results for the variance decomposition analysis to adaptation (MAD) and alternate error reduction strategies (AERS), and all associated post-hoc tests for day 1 and day 2. Associations at *p*_*unc*_ < 0.05 are bolded. In both tables, REST1 has been abbreviated as R1, REST2*−*REST1 as R2-1, and REST3*−*REST1 as R3-1.

| ROI Pair | Behavior | p(model) | Constant | R1 | p(R1) | R2-1 | p(R2-1) | R3-1 | p(R3-1) | D1 | p(D1) |
| --- | --- | --- | --- | --- | --- | --- | --- | --- | --- | --- | --- |
| Left M1-Left S1 | D1 RER | 0.0088 | 0.51 | 0.24 | **0.0088** | - | - | - | - | - | - |
| Left S1-Right M1 | D1 RER | **0.0007** | 0.59 | - | - | - | - | -0.34 | **0.0007** | - | - |
| Left PM-Right M1 | D1 MAE | **0.0033** | 13.44 | 9.85 | 0.0480 | -10.68 | 0.0567 | 19.82 | **0.0006** | - | - |
| Left M1-Right CB6 | D1 MAE | 0.0100 | 16.82 | - | - | 20.05 | **0.0097** | -16.19 | **0.0146** | - | - |
| Right CB6-Right CB8 | D1 MAI | 0.0089 | 0.35 | - | - | - | - | 0.48 | **0.0089** | - | - |
| Left M1-Right CB8 | D2 RER | **0.0005** | 0.2 | - | - | - | - | 0.42 | **0.0012** | 0.64 | 0.0025 |
| Left PM-Right CB8 | D2 RER | **0.0008** | 0.34 | -0.42 | **0.0020** | - | - | - | - | 0.49 | 0.0161 |
| Left PM-Right CB8 | D2 MAE | 0.0083 | 14.32 | - | - | - | - | 17.1 | **0.0037** | 0.19 | 0.2853 |
| Left M1-Right CB6 | D2 MAI | **0.0030** | 0.22 | 0.58 | **0.0027** | 0.4 | 0.0508 | - | - | 0.43 | 0.0339 |
| Left S1-Right CB6 | D2 MAI | **0.0013** | 0.15 | 0.47 | **0.0117** | 0.69 | **0.0014** | - | - | 0.53 | 0.0083 |
| Right CB6-Right CB8 | D2 MAI | 0.0074 | 0.08 | 0.41 | 0.0290 | - | - | 0.63 | **0.0160** | 0.38 | 0.0160 |

**Table 1.** Top: Results of model based analysis of rsFC to behavior (RER, MAE, MAI) measured on day 1 and day 2.
| ROI Pair | rsFC | p(model) | Constant | D1 AERS | D1 MAD | p(AERS) | p(MAD) | Post-hoc Direct Associations |
| --- | --- | --- | --- | --- | --- | --- | --- | --- |
| Left M1-Left S1 | R1 | 0.026 | 0.14 | 0.74 | 0.26 | **0.010** | 0.401 | D1 AERS ∼ 3.62 + 2.78(R1), $p < 0.0097$ |
| Left S1-Right M1 | R3-1 | 0.002 | 0.24 | -0.80 | -0.27 | **0.001** | 0.276 | D1 AERS ∼ 0.28 - 0.42(R3-1), $p < 0.0007$ |
| Right CB6-Right CB8 | R3-1 | 0.045 | -0.09 | 0.10 | 0.44 | 0.500 | **0.017** | D1 MAD ∼ 0.25 + 0.43(R3-1), $p < 0.0158$ |
| Left PM-Right M1 | R1 | 0.172 | 0.20 | 0.47 | 0.23 | 0.084 | 0.467 | D1 AERS ∼ 0.23 + 0.22(R1), $p < 0.0814$ |
| Left PM-Right M1 | R3-1 | 0.074 | -0.01 | -0.59 | 0.22 | **0.031** | 0.459 | D1 AERS ∼ 0.29 - 0.27(R3-1), $p < 0.029$ |
| Left PM-Right M1 | R3-2 | 0.158 | -0.05 | -0.34 | 0.44 | 0.196 | 0.150 | D1 MAD ∼ 0.28 + 0.16(R3-2), $p < 0.1550$ |
| Left M1-Right CB6 | R2-1 | 0.277 | 0.03 | -0.19 | 0.26 | 0.297 | 0.222 | D1 MAD ∼ 0.27 + 0.20(R2-1), $p < 0.2227$ |
| Left M1-Right CB6 | R3-1 | 0.736 | 0.18 | -0.09 | -0.17 | 0.683 | 0.508 | D1 MAD ∼ 0.29 - 0.10(R3-1), $p < 0.5019$ |
| Left M1-Right CB6 | R3-2 | 0.251 | 0.15 | 0.10 | -0.43 | 0.654 | **0.111** | D1 MAD ∼ 0.29 - 0.21(R3-2), $p < 0.1061$ |
| ROI Pair | rsFC | p(model) | $\beta_0$ | D2 AERS | D2 MAD | p(AERS) | p(MAD) | Post-hoc Direct Associations |
| Left PM-Right CB8 | R1 | 0.013 | 0.40 | -0.45 | -0.28 | **0.010** | 0.122 | D2 AERS ∼ 0.32 + -0.47(R1) + 0.24(D1), $p < 0.022$; $p(D1) < 0.2610$, $p(R1) < 0.010$ |
| Left M1-Right CB8 | R3-1 | 0.006 | -0.33 | 0.26 | 0.59 | 0.140 | **0.004** | D2 MAD ∼ 0.34 + 0.42(R3-1) + 0.19(D1), $p < 0.016$; $p(D1) < 0.409$, $p(R3-1) < 0.009$ |
| Left PM-Right CB8 | R3-1 | 0.004 | -0.33 | 0.30 | 0.56 | 0.073 | **0.004** | D2 MAD ∼ 0.32 + 0.43(R3-1) + 0.28(D1), $p < 0.008$; $p(D1) < 0.206$, $p(R3-1) < 0.005$ |
| Left M1-Right CB6 | R1 | 0.062 | -0.05 | -0.17 | 0.36 | 0.268 | **0.035** | D2 MAD ∼ 0.28 + 0.41(R1) + 0.31(D1), $p < 0.047$; $p(D1) < 0.186$, $p(R1) < 0.034$ |
| Left M1-Right CB6 | R2-1 | 0.673 | -0.02 | 0.02 | 0.14 | 0.907 | 0.382 | D2 MAD ∼ 0.30 + 0.14(R2-1) + 0.27(D1), $p < 0.385$; $p(D1) < 0.293$, $p(R2-1) < 0.529$ |
| Left M1-Right CB6 | Full model: D2 MAD ∼ 0.29 + 0.57(R1) + 0.42(R2-1) + 0.20(D1), $p < 0.0246$; $p(R1) < 0.0071$, $p(R2-1) < 0.0705$, $p(D1) < 0.3770$ | | | | | | | |
| Left S1-Right CB6 | R1 | 0.682 | 0.08 | -0.14 | 0.02 | 0.391 | 0.902 | D2 MAD ∼ 0.30 + -0.01(R1) + 0.31(D1), $p < 0.471$; $p(D1) < 0.226$, $p(R1) < 0.970$ |
| Left S1-Right CB6 | R2-1 | 0.269 | -0.01 | 0.02 | 0.32 | 0.911 | 0.109 | D2 MAD ∼ 0.28 + 0.34(R2-1) + 0.32(D1), $p < 0.141$; $p(D1) < 0.191$, $p(R2-1) < 0.124$ |
| Left S1-Right CB6 | Full model: D2 MAD ∼ 0.27 + 0.18(R1) + 0.43(R2-1) + 0.30(D1), $p < 0.2147$; $p(R1) < 0.4356$, $p(R2-1) < 0.0894$, $p(D1) < 0.2353$ | | | | | | | |
| Right CB6-Right CB8 | R1 | 0.512 | 0.37 | -0.17 | 0.13 | 0.345 | 0.507 | D2 AERS ∼ 0.32 + -0.21(R1) + 0.24(D1), $p < 0.382$; $p(D1) < 0.314$, $p(R1) < 0.318$ |
| Right CB6-Right CB8 | R3-1 | 0.186 | -0.05 | 0.20 | 0.14 | 0.124 | 0.311 | D2 AERS ∼ 0.24 + 0.39(R3-1) + 0.19(D1), $p < 0.225$; $p(D1) < 0.419$, $p(R3-1) < 0.154$ |
| Right CB6-Right CB8 | Full model: D2 AERS ∼ 0.27 + -0.09(R1) + 0.33(R3-1) + 0.20(D1), $p < 0.3769$; $p(D1) < 0.4041$, $p(R1) < 0.6856$, $p(R3-1) < 0.2809$ | | | | | | | |

Following the dynamic perturbation task (REST3 − REST1), interhemispheric decreases in rsFC between cortical sensori-motor areas were significantly associated with day 1 relative error reduction and magnitude of after effects. Change in interhemispheric rsFC was negatively associated with relative error reduction (RER *∼* L-S1 to R-M1, *p*_*model*_ = 0.0007, *p*_*REST* 3*−REST* 1_ = 0.0007), and positively associated with magnitude of after effects (MAE *∼* L-PM to R-M1, *p*_*model*_ =0.0033, *p*_*REST*3*−REST*1_ = 0.0006). For left PM to right M1, these effects were more significant across the dynamic perturbation task (REST3 − REST2), indicated by a post-hoc test conducted on the reduced model (reduced model contrast: *p*_*REST* 3*−REST* 2_ = 0.0001, D1 MAE = 16.48+ 15.50(REST3 − REST2), *p* = 0.0021). Post-hoc analysis showed that these associations were driven by decreases in rsFC within these edges correlating with greater relative error reduction and lower magnitude of after effects. In line with these results, decreases in rsFC in both edges was preferentially associated with alternate error reduction strategies (AERS ∼ L-S1 to R-M1, *p* = 0.001; AERS ∼ L-PM to R-M1, *p* = 0.031). These results suggest that decreases in rsFC between these nodes reflect a preferential use of alternate error reduction strategies rather than adaptation, that results in performance error reduction without significant after effects.

#### Associations between behavior and rsFC in the cortico-thalamic-cerebellar network

Within the cortico-thalamic-cerebellar network, there were no significant associations between baseline rsFC (REST1) and day 1 behavior. Linear regression analysis to day 2 behavior returned significant (*p*_*FWE*_ < 0.05) associations to baseline rsFC in three edges (Tbl. 1). Baseline rsFC between the left sensorimotor cortex (L-M1, L-S1) and the anterior cerebellum (R-CB6) was positively associated with magnitude of adaptation (MAI ∼ L-M1 to R-CB6, *p*_*model*_ = 0.0030, *p*_*REST*1_ = 0.0027; MAI ∼ L-S1 to R-CB6, *p*_*model*_ = 0.0013, *p*_*REST*1_ = 0.012), while lower baseline rsFC between the left sensorimotor cortex (L-PM) and the posterior cerebellum (R-CB8) was associated with greater relative error reduction (RER, *p*_*model*_ = 0.0008, *p*_*REST* 1_ = 0.002). Variance decomposition analysis showed that baseline rsFC between the left motor cortex to right anterior cerebellum was preferentially associated with adaptation (*p* = 0.035), while lower baseline rsFC between the left premotor cortex and the right posterior cerebellum was associated with a greater reliance on alternate error reduction strategies (*p* = 0.004, Tbl. 1). For other nodes in this network (L-S1 to R-CB6) there was no clear association between baseline rsFC and either control strategy.

Linear regression analysis returned a limited set of associations between change in rsFC following the motor performance task (REST2 − REST1) and behavior. For day 1, no model reached significance when correcting for multiple comparisons. However, at *p*_*unc*_ < 0.01, change in rsFC between the left motor cortex to the right anterior cerebellum was positively associated with after effects (MAE *∼* L-M1 to R-CB6, *p*_*model*_ = 0.010, *p*_*REST* 2*−REST* 1_ = 0.0097, Tbl. 1). For Day 2, an edge in the same network showed a significant (*p*_*FWE*_ < 0.05) positive association to magnitude of adaptation at day 2 when correcting for day 1 behavior (MAI *∼* L-S1 to R-CB6, *p*_*model*_ = 0.0013, *p*_*REST* 2*−REST* 1_ = 0.0014, Tbl. 1). The variance decomposition analysis showed that these changes were preferentially associated with adaptation, though no significant associations were identified (MAD ∼ L-M1 to R-CB6, *p* = 0.071; MAD ∼ L-S1 to R-CB6, *p* = 0.089).

Linear regression analysis returned a larger set of associations between change in rsFC following the dynamic perturbation task (REST3 − REST1) and behavior. For day 1, no model reached significance when correcting for multiple comparisons. At *p*_*unc*_ < 0.01, decreases in rsFC between the motor cortex to the anterior cerebellum were associated with greater magnitude of after effects (MAE *∼* L-M1 to R-CB6, *p*_*model*_ = 0.01, *p*_*REST* 3*−REST* 1_ = 0.0146). Post-hoc testing conducted on the reduced model showed that this association reached significance (*p*_*FWE*_ < 0.05) across the dynamic perturbation task (reduced model contrast: *p*_*REST* 3*−REST* 2_ = 0.0027, D1 MAE = 17.08 -17.69(REST3 − REST2), *p <* 0.0026). Variance decomposition analysis returned a weak association between decreases in rsFC between these nodes and adaptation specific processes (*p* = 0.11). Additionally, there was a positive association between increases in rsFC within the cerebellum and adaptation (MAI *∼* R-CB6 to R-CB8, *p*_*model*_ = 0.0089, *p*_*REST* 3*−REST* 1_ = 0.0089), that was attributed to adaptation specific processes (MAD ∼ R-CB6 to R-CB8, *p* = 0.017).

For analysis of day 2 behavior, increases in rsFC within the cortico-thalamic-cerebellar network were positively associated with greater adaptation on day 2 when controlling for day 1 behavior (Tbl. 1). Increases in rsFC between left motor cortex (L-M1) and the posterior cerebellum (R-CB8) were significantly associated (*p*_*FWE*_ < 0.05) to gains in relative error reduction (RER, *p*_*model*_ = 0.0005, *p*_*REST* 3*−REST* 1_ = 0.0012). Post-hoc testing showed similar positive associations to gains in magnitude of adaptation (D2 MAE = 14.53 + 16.42(REST3*−*REST1) + 0.17(D1 MAE), *p*_*model*_ = 0.017, *p*_*REST*3*−REST*1_ = 0.0083; D2 MAI = 0.28 + 0.31(REST3*−*REST1) + 0.38(D1 MAI), *p*_*model*_ = 0.016, *p*_*REST* 3*−REST* 1_ = 0.048). Edges identified without full-network FWE correction (*p*_*unc*_ < 0.01) included the L-PM to R-CB8 (MAE, *p*_*model*_ = 0.0083, *p*_*REST*3*−REST*1_ = 0.004), and R-CB6 to R-CB8 (MAI, *p*_*model*_ = 0.0074, *p*_*REST*3*−REST*1_ = 0.016). The variance decomposition analysis identified a significant association between the increases in rsFC in the cortico-cerebellar edges and adaptation (MAD ∼ L-M1 to R-CB8, *p* = 0.004; MAD ∼ L-PM to R-CB8, *p* = 0.004). For the intra-cerebellar edge (R-CB6 to R-CB8), analysis to day 2 behavior returned a reduced model that identified an association between both baseline rsFC and change in rsFC after the dynamic perturbation task (Tbl. 1). Change in rsFC between the REST3 − REST1 conditions and day 1 behavior were significantly associated, which can confound the linear regression analysis of rsFC to day 2 behavior. As such, we performed a post-hoc analysis of change in magnitude of adaptation to rsFC. Baseline rsFC remained positively associated with greater gains in adaptation, (ΔMAI = −0.10 + 0.37(REST1), *p* = 0.045), however increases in rsFC were not significantly associated to changes in behavior (ΔMAI = 0.03 + 0.09(REST3 − REST1), *p* = 0.719). Variance decomposition analysis returned no significant associations to either alternate error reduction strategies nor adaptation for rsFC measured in this edge.

## Discussion

The purpose of this study was to identify the neural processes associated with motor memory formation that are specific to dynamic adaptation of the wrist immediately following task performance. Using an fMRI-compatible wrist robot, the MR-SoftWrist, participants performed wrist pointing under dynamic perturbations. Task-based fMRI was used to localize brain regions associated with active motor control processes, which included a sensorimotor network and the cortico-thalamic-cerebellar network of the trained right wrist. We quantified changes in resting-state functional connectivity (rsFC) within these networks immediately following task performance. To determine associations between rsFC measured in these networks, adaptation behavior, and motor memory formation, we performed a linear regression between rsFC and behavioral metrics measured on day 1, and on day 2 while controlling for day 1 behavior. To identify networks associated specifically with adaptation, we performed post-hoc linear regression of rsFC measured in these networks to behavioral metrics attributed to adaptation or alternate error reduction strategies. Critically, our work provides new insight on how rsFC in the sensorimotor and cortico-thalamic-cerebellar networks is modulated immediately after adaptation to dynamic perturbations, and on how these modulations are associated to adaptation-specific behavior and its retention.

### Behavior

Behavioral analysis showed significant evidence of error reduction across the dynamic perturbation task, indicative of motor learning. Adaptation-specific features, including significant after effects and increases in adaptation index, suggest that participants formed an internal model of the lateral force environment to reject perturbations applied to the wrist. Given the weak correlation between relative error reduction and the magnitude of adaptation index 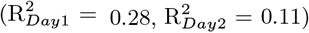 and magnitude of after effects 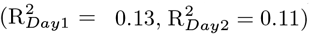, alternate error reduction strategies—such as increased joint impedance via muscle co-contraction (2)— likely contributed to observed behavior. Using a variance decomposition analysis, we identified independent components of behavior specific to adaptation and to alternate error reduction strategies. In our behavioral data, alternate error reduction strategies were shown to account for 72-88% of the observed variance in relative error reduction, while adaptation accounted for just 26-12% (Day 1–Day 2, respectively). Moreover, the end magnitude of adaptation achieved in our study (mean ± s.e.m.: 0.355 ± 0.0257) was lower than is commonly reported in the literature for dynamic adaptation in reaching or visuomotor adaptation following a similar number of trials (0.5 < adaptation index < 0.85) (23–25). As the intrinsic dynamics of the wrist joint are dominated by stiffness, this discrepancy could be due to a difference in relative neuromotor control mechanisms utilized during task performance, such as a greater reliance on alternate error reduction strategies in the wrist compared to reaching tasks (26, 27).

Between days, there was behavioral evidence of retention, including significant decreases in initial errors (*p* = 0.0103), consistent with recall processes (20), significantly lower final errors (*p* = 0.044) and an increase in end magnitude of adaptation (*p* = 0.056) on Day 2 compared to Day 1. There was no significant change in relearning at the group level, as is commonly reported in the adaptation literature (20, 28, 29). The performance of the resistive force task between days may have washed out these effects in our study (Fig. 1), and contributed to the weaker effects of retention observed.

### Changes in rsFC

Within the sensorimotor network, rsFC decreased interhemispherically following both the motor performance task and the dynamic perturbation task. Following the motor performance task, changes in rsFC were localized to the motor cortices (L-PM to R-M1, L-PM to R-S1) and between the primary motor cortex and the left thalamus. The motor cortex is engaged in motor task performance in the absence of learning, and has significant projections to the thalamus (30). Given the immediate acquisition of resting state data following task performance, these changes may reflect residual stimulation of the motor network following task performance. Moreover, these effects likely have a small effect size, as they did not reach significance in either of the supplementary analyses conducted (i.e., Expanded Network Analysis, ROI-to-Voxel Analysis). Following the dynamic perturbation task, the interhemispheric decreases remained significant when correcting for rsFC measured in REST2, and an additional decrease between the L-M1 and R-M1 nodes was observed. Moreover, both supplemental analyses returned significant interhemispheric decreases for the REST3 − REST1 contrasts, but not for REST2 − REST1. Previously, a decrease in rsFC between the motor cortices has been measured immediately following a robot-mediated upper extremity reaching task and was associated with motor learning in stroke patients (21). However, these decreases have not been observed in neurotypical adults. Decreases in rsFC may reflect neural processes occurring within the trained left motor cortex that are independent from the right motor cortex, in line with an inhibitory connection between the two motor cortices and an increased segregation of cross-cortical activity (31).

Within the cortico-thalamic-cerebellar network, no changes were observed with motor execution. Following the dynamic perturbation task, increases in rsFC between the left cortical sensorimotor regions to both the left thalamus and the right cerebellum were observed across all analysis modalities. These effects are in line with previous works that have identified change in rsFC in this network following adaptation (16, 18, 20). However, previous studies report an inhibitory (decreases in rsFC) relationship between motor and cerebellar nodes following adaptation, that differs from the facilitory (increases in rsFC) relationship observed in our study. TMS studies conducted during visuomotor adaptation have shown a faciliatory relationship between the cerebellum and the motor cortex early in the adaptation process, when errors are large, and an inhibitory pattern (typically observed at rest) that returned following acquisition of the learned motor pattern when errors are small (32). Given the immediate acquisition of our resting state scans, our effects may capture a transient faciliatory relationship between the cerebellum and the primary cortex that has not been observed in rsFC data taken at longer timescales. Moreover, previous studies that reported an inhibitory relationship achieved a greater degree of error reduction at the group level that may contribute to the different pattern of rsFC observed (16).

### Associations between rsFC and behavior

RsFC measured within the sensorimotor and cortico-thalamic-cerebellar networks was associated with different motor control strategies.

Within the sensorimotor network, rsFC was only associated with day 1 behavior that was attributed to alternate error reduction strategies. Greater baseline rsFC within the left motor cortex was positively associated with greater relative error reduction, while interhemispheric decreases in rsFC following the dynamic perturbation task were associated with greater relative error reduction and lower magnitude of after effects. These results suggests that engagement of the motor network is associated with a greater reliance on alternate motor control processes instead of adaptation to reduce performance errors. Recently, decreases in rsFC between the left M1 and right sensorimotor areas were associated with initial increases in co-contraction in a dynamic perturbation task (18). Taken together, these results suggest that immediate changes in rsFC in the sensorimotor network following task performance reflect neural processes associated with alternate motor control strategies, potentially co-contraction, that are independent from adaptation processes.

Instead, within the trained cortico-thalamic-cerebellar network, rsFC was associated with adaptation specific processes, and retention of adaptation between days. Between the left motor cortex and right anterior cerebellum (R-CB6), greater baseline and pre-task rsFC (REST1, REST2 − REST1) was associated with greater adaptation on day 1 (MAE), and gains in adaptation (MAI) on day 2. Baseline effects were preferentially associated with adaptation. Following performance of the dynamic perturbation task, decreases in rsFC between the left M1 and right CB6 were associated with greater adaptation (MAI), consistent with previous findings in the literature (16). In contrast, increases in rsFC between the left motor cortex (L-M1, L-PM) and right posterior cerebellum (R-CB8) were positively associated with greater relative error reduction, and with increases in our adaptation metrics (MAI, MAE), that were attributed to adaptation specific processes in our variance decomposition analysis. These results highlight a differential pattern of connectivity between the motor cortex and the posterior cerebellum (increases) and the anterior cerebellum (decreases) following adaptation. The differential patterns of activation between the anterior and posterior cerebellum could reflect their different roles in adaptation learning, as the posterior cerebellum has been associated with fast adaptive states that are highly reactive to large errors, while the anterior cerebellum has been associated with slow adaptation states that peak when errors are small (25, 33), although this was not directly investigated in our study. Finally, rsFC measured within the cerebellum showed a weak (*p*_*unc*_ *<* 0.01) positive association to adaptive behavior. Increases in rsFC following the dynamic perturbation task were associated with greater adaptation on day 1, and baseline rsFC was positively associated with gains in adaptation on day 2. These results agree with previous findings that the cerebellum is engaged in active adaptation, but not necessarily in memory formation (9). Overall, these results agree with previous findings that the cortico-cerebellar network is integral to dynamic adaptation (11, 34, 35).

In summary, our study supports the hypothesis that changes in rsFC measured immediately after adaptation are reflective of motor memory formation processes, and provides evidence that different brain networks—the sensorimotor network and the trained cortico-thalamic-cerebellar network—are involved in different motor control processes that contribute to improvement in task performance in response to dynamic perturbations of the wrist.

## Materials and Methods

Thirty four right-handed young adults (21 M, 13 F, age: 24 ± 4 years), free from neurological or musculoskeletal injury participated in this study. Two of these participants were excluded due to hardware failures (F, 28 years old; F, 30 years old), and two participants were excluded for abnormal task performance (M, 21 years old; M, 25 years old). The study was approved by the Institutional Review Board of the University of Delaware, IRB no 906215-10.

### Experimental Protocol

The experimental protocol is shown in Fig. 1, A. On day 1, participants first performed a training task in the zero force condition to familiarize them with the MR-SoftWrist and task instructions. After training, participant and device set-up in the MRI took roughly 30 minutes. The fMRI protocol began with 10 minutes of structural scans, followed by three resting states scans and three motor tasks. All motor tasks followed a blocked design (Fig. 1, A), with 15 s rest blocks interleaved between blocks of 24 active trials. During rest blocks, the MR-SoftWrist held the participants’ hand stationary while a virtual cursor was displayed performing straight point-to-point movements between the targets to match the visual stimulus seen during task performance.

The first resting state scan (REST1) was taken to assess baseline resting state functional connectivity. During all resting state scans, participants were instructed to fixate on a white cross displayed on a black background. Participants then performed the motor performance task, which consisted of 144 trials (6 blocks) in a zero force condition. REST2 was acquired immediately following task completion to measure neural activity at rest following normal execution of wrist pointing. Following REST2, participants performed a dynamic perturbation task that included 168 trials (7 blocks). Blocks 1-6 were performed in a velocity dependent lateral force condition (6), and the last block was performed in the zero force condition to enable assessment of after effects. REST3 was acquired immediately following the dynamic perturbation task to measure neural activity at rest following training of a new motor control pattern. Subjects then performed a resistive force task, which consisted of 168 trials (7 blocks). The first block was performed in the zero force condition, and blocks 2-7 were performed in the resistive force condition. For the purposes of this study we only analyzed data from the first block of this task.

On day two, participants performed a behavioral follow-up task in a mock scanner to assess retention of day one training. The mock scanner was built to the same dimensions as the MR-scanner and allowed participants to perform the follow-up task in the same supine position as day one. The follow up task consisted of 240 trials (10 blocks) with 15 s rest conditions interleaved. The first (1-2) and last (9-10) two blocks were performed in the zero force condition, while blocks 3-8 were performed in the lateral force condition.

### MR-SoftWrist

The MR-SoftWrist is an fMRI-compatible wrist robot (Fig. 1, B) that supports wrist pointing in flexion-extension (FE) and radial-ulnar deviation (RUD) in a circular workspace (radius = 20 deg). The device has a maximum output torque of 1.5 Nm about each axis and implements gravity compensation and force feedback at 1000 Hz. The MR-SoftWrist can display a variety of kinesthetic environments to the user, ranging from a zero force mode where it minimally perturbs the user’s movements (peak resistive torque < 0.17 Nm, trial average resistive torque = 0.05 Nm), to a stiffness control mode where it displays a high stiffness environment (max virtual stiffness *k*_*v*_ =0.45 Nm/deg) (3). The MR-SoftWrist introduces no change in the noise characteristics of functional images or measures of neural activity acquired during participant interaction with the robot (36). For details on the control and design of the MR-SoftWrist, see (3).

### Task Design

For all motor tasks, participants grasped the handle of the MR-SoftWrist and moved their wrist to control a grey cursor (radius 1 deg) continuously displayed on a monitor. Flexion-extension of the wrist moved the cursor horizontally, while radial-ulnar deviation moved the cursor vertically. Pronation-supination was prevented by a forearm support. Subjects were cued to make alternating flexion-extension rotations to move the cursor in a straight line to one of two targets (radii 1.25 deg) located at (±10, 0) degrees in flexion-extension, radial-ulnar deviation respectively (Fig. 1, B). Trial onset was cued by a change in target color from black to blue. Trial completion was achieved when the error between the cursor and the target was less than 2 degrees for more than 250 ms. After trial completion, the reached target provided timing feedback for 0.5 s by turning red if the movement duration was greater than 650 ms or green if it was less than 300 ms. Otherwise, the target remained black. Duration of the inter-trial interval between feedback and cuing of the next trial was chosen at random from a truncated normal distribution with a mean ± standard deviation of 0.75 ± 0.25 s, bounded by [0.25 1.75] s.

The robot operated in one of four control modes: a zero force mode (ZF), a lateral force mode (LF), a resistive force mode (RF), and an error clamp mode (EC) (Fig. 1, C). In the ZF mode, the desired interaction torque was set to zero (*τ*_*FE*_ = 0; *τ*_*RUD*_ = 0) to enable measurement of wrist-pointing behavior in a transparent environment. In the LF mode, the robot applied a velocity-dependent torque proportional to the measured velocity as [*τ*_*FE*_; *τ*_*RUD*_] = [0 *B*; − *B* 0] [*v*_*FE*_; *v*_*RUD*_]), B = 14.32 Nm·s/deg, which creates a clockwise force field that applies forces perpendicular to the direction of movement (6). In the RF mode, the robot applied a velocity-dependent torque proportional to the measured velocity as [*τ*_*FE*_; *τ*_*RUD*_] = [− *B* 0; 0 −*B*] [*v*_*FE*_; *v*_*RUD*_], *B* = 14.32 N·ms/deg, resulting in a resistive force applied opposite to the direction of movement. In the error clamp mode, the robot applied a force channel (deadband = 0.1 deg, stiffness = 15 N·m/deg, damping = 34.4 N·ms/deg) that restricted lateral deviations to < 0.5 deg, to enable measurement of lateral force profiles generated by participants that reflect their expectation of required task dynamics (19). Error clamp trials were interspersed throughout the ZF, LF, and RF conditions with a 1/8 probability.

### Behavioral Data Analysis

All behavioral data were processed using MATLAB (The MathWorks, version 2020b). Velocity and force data were recorded with a sampling rate of 1000 Hz, and position data was recorded with a sampling rate of 100 Hz. For data processing, position data were up-sampled to a 1000Hz sampling rate using the resample function. All position, velocity, and force data were then low-pass filtered at 25 Hz with a 4th order Butterworth filter applied via the filtfilt function. For trial-by-trial data analysis, trial onset was defined as the instant the absolute cursor velocity exceeded 20 deg/s. Trial end was defined as the instant the cursor was within 3 degrees of the target in the flexion-extension direction. Trials were excluded if they matched any of the following conditions: 1) a trial duration of less than 200 ms, or greater than 700 ms; 2) a maximum velocity below 40 deg/s; 3) a reversal in goal directed velocity occurring before the trial max velocity, indicative of false starts. At the group level, this resulted in an exclusion of 3.1 - 2.8% (day 1 - day 2) of all field trials (zero-force, lateral force) and 1.8 - 2.8% of error clamp trials (day 1 - day 2, respectively).

For each trial, position and force data from trial onset to trial end were upsampled into 1000 data points and divided between extension and flexion movements. For day 1, baseline trajectory and force profiles in each direction were calculated as the average of all valid right (extension) and left (flexion) directed trials in blocks 3-6 of the motor performance task. For day 2, baseline trajectories were determined as the average profile of all valid trials performed in the zero force condition in the first two blocks of the behavioral follow up task. Within each day, direction-specific baseline profiles were subtracted from the respective right- or left-directed trials for force and position data, such that all behavioral metrics reported reflect a change relative to typical, non-perturbed wrist pointing.

#### Behavioral metrics

Trajectory errors were calculated in all conditions as the internal angle between the cursor and the start and end targets, taken at the cursor’s maximum lateral deviation within the first 150 ms after trial onset. To quantify wrist kinetics, we calculated participants’ adaptation index as the area under the force profile measured on error clamp trials divided by the area under the ideal force profile necessary to compensate for the force field, within the first 150ms after trial onset. As error clamp trials occur unexpectedly, force profiles measured in these trials reflect participants expectation of the required task dynamics. An adaptation index of one represents perfect adaptation, while an adaptation index of zero represents no adaptation. By restricting our behavioral metrics to the first 150 ms of the trial, we aimed to capture primarily feed-forward motor control processes that most directly reflect pre-planned motor actions (37).

#### Group Level Behavioral Analysis

We used two repeated-measures one-way ANOVAs to test for an effect of experimental phase on trajectory error and adaptation index measured across tasks performed on both days. For adaptation index, the experimental phases of interest for day one included Baseline, defined for each participant as the average adaptation index measured in the last block of the motor performance task, Early LF and Late LF, defined as the average adaptation index measured in the first and the last block of trials performed in the lateral force condition, respectively, and After Effects, defined as the average adaptation index measured in the block of zero-force trials immediately following the lateral force condition in the dynamic perturbation task (Fig. 1, bottom left). For day two, experimental phases were defined similarly, such that day two Baseline, Early LF, Late LF and After Effects phases corresponded to the average adaptation index measured in blocks 2, 3, 8 and 9 of the behavioral follow up task, respectively.

For trajectory error, the Baseline and Late LF experimental phases were defined consistently with the phases used for adaptation index data. To investigate two aspects of retention reported in the literature—recall and relearning—we divided the Early LF phase into two parts (20). Recall refers to decreases in initial errors between days, while relearning refers to increases in the rate of error reduction in the subsequent trials between days. We defined Initial Error phases as the average trajectory error measured in trials 2-4 of block 1 (lateral force condition) of the dynamic perturbation task for day one, and block 3 of the behavioral follow up task for day two. The first trial was excluded as it is assumed to act as the initial cue to participants to recall any previously learned response to the lateral force field. Middle Errors were defined as the average trajectory error measured in the subsequent trials (5-24) in block 1 and block 3, respectively. Finally, the After Effects phase was defined as the average trajectory error measured in trails 1-4 of the no force blocks performed immediately following the lateral force condition on each day, as after effects in trajectory error data are highly transient.

When the ANOVA returned a significant effect, post-hoc Tukey test were used to quantify the effect of experimental phase on adaptation index and trajectory error. Contrasts between parameter estimates of experimental phases between days were used to determine effects of retention.

#### Behavioral Variance Decomposition

To isolate behavioral effects associated specifically with adaptation from those associated with alternate error reduction strategies, we performed a two-part variance decomposition analysis using our behavioral summary metrics (RER, MAE, and MAI). We used the following set of equations:

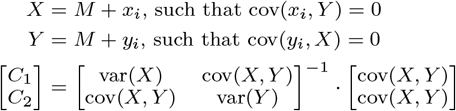

to identify variance that is mutually attributable to both metrics of interest (*M* = *C*_1_ ·*X* + *C*_2_ ·*Y*), and variance that is attributable to a single metric of interest (*x*_*i*_ or *y*_*i*_) that is uncorrelated with the other metric (*Y* or *X*, respectively) (16).

First, we found a mutual component of adaptation (MAD) from our two adaptation specific metrics (MAE and MAI), in which *X* = MAE, *Y* = MAI, and *M* = MAD. This analysis returned an individual estimate of MAD for each subject. Next, we performed a variance decomposition using the resultant MAD metric paired with RER to isolate variation within the RER metric that is independent from adaptation processes. Here, *X* = RER, *Y* = MAD, and *x*_*i*_, the component of RER that is independent from MAD, was used to quantify alternate error reduction strategies (AERS) from adaptation processes. These two independent metrics were then used in an exploratory analysis to determine if associations between rsFC and behavior were driven by internal model formation processes (MAD), or alternate error reduction strategies (AERS).

### fMRI Data Analysis

All fMRI data were acquired at the Center for Biological and Biomedical Imaging at the University of Delaware, on a 3.0 T Siemens Prisma MR scanner using a 64-channel head coil. Subjects laid supine in the scanner with their head immobilized by foam padding. Earplugs were provided to reduce scanner noise, as were headphones for communication with experimenters in between imaging sequences. Following a brief initial localizer scan (Siemens AA Head Scout, spatial resolution: 1.6×1.6×1.6 mm^3^, isotropic; TR = 3.15 s, TE = 1.37 ms), we performed a T1-weighted anatomical scan (spatial resolution 1×1×1 mm^3^, isotropic; TE = 3.02 ms; TR = 2.30 s), followed by a GRE field map (spatial resolution 3×3×3 mm^3^, TR = 400 ms, TE = 4.92 ms). All subsequent functional scans (task-based and resting state) were acquired using a multi-band T2^*∗*^-weighted EPI sequence (axial slices oriented to the AC-PC line, spatial resolution 2×2×2 mm^3^, 60 slices, TE = 30 ms, TR = 1 s, acceleration factor = 4).

#### fMRI Data Pre-processing

Pre-processing of fMRI data was performed using the SPM12 software (38). All task-based and resting state functional images were unwarped and field map corrected, realigned, normalized into standard (MNI) space for group analysis, and smoothed with a 6 mm gaussian kernel. We used the artifact detection toolbox (ART) to identify outlier volumes, defined as volumes with frame-wise displacement above 0.9 mm or global BOLD signal changes above 5 standard deviations (39). Outlier scans, along with rotational and translation head movement parameters were included as nuisance regressors of non-interest in our tasked based analysis. For resting-state data, we used a multiple regression analysis to remove effects of non-interests, that included the same nuisance regressors as our task-based data as well as the first 5 principal components extracted from the white matter and CSF tissue maps to model physiological noise (40). Resting state data were bandpass filtered between 0.008–0.09 Hz, as signal related to neural activity is expected to lie within this frequency range (41).

#### ROI identification

Task-based fMRI scans were used to determine relevant regions of interest (ROIs) for analysis of our resting state data. For all motor tasks, we created task-related regressors using a boxcar function to model active blocks and rest blocks, that were then convolved with the standard hemodynamic response function. Regressors were entered into a first-level GLM analysis, and contrasts between coefficients associated with task regressors and rest were used to produce subject-specific t-maps of task-related activation. For the dynamic perturbation task, a second-level one-sample t-test was used to identify group-level activations associated with dynamic motor control. Using this map (thresholded at *t >* 5.63, derived from a *p*_*FDR*_ *<* 0.001 voxel-level correction), we identified all anatomical regions with significant activation. We partitioned these task-related regions into a total of 65 anatomical ROIs, using the Juelich histological atlas for cortical regions (42), the Talaraich atlas for the frontal poles, the Harvard-Oxford atlas for sub-cortical regions (43), and the MNI FLIRT atlas for cerebellar regions (44), all thresholded at 50% probability.

For the left cortical areas in the sensorimotor network, which are highly engaged during task execution, we defined subject-specific ROIs based on task related activation measured within each anatomical sensorimotor ROI. Subject specific t-maps corresponding to the contrast of lateral force > rest were thresholded at *t >* 0 and weighted by the contrast of lateral force > rest – zero force > rest. Using these maps, we constructed a spheroid centered on the center of mass of task related activation within each anatomical ROI, using the fslmaths function (https://fsl.fmrib.ox.ac.uk/). ROIs were created with a target volume of 180 voxels, which corresponds to a spheroid with a radius of 7 mm (20). However, as all ROIs were bounded by the borders of the anatomical ROI, the spheroids radii were iteratively increased to reach the desired volume. Spheroid radii were increased by 1 mm up to a max 14 mm, as long as the absolute error between the desired and achieved volume decreased and the activation cluster within the ROI remained contiguous.

#### Changes in resting state functional connectivity

All resting state fMRI analyses were performed using the CONN toolbox (45). At the subject-level, the timeseries of each ROI was calculated as the average timeseries across all voxels within the ROI for each resting state condition. For all ROI-based analysis, resting state functional connectivity (rsFC) between each ROI pair was quantified by a scalar, as the Fisher-transformed correlation coefficient between the timeseries of each ROI, in each condition. For ROI-to-voxel level analysis, rsFC was quantified using the ROI timeseries as a regressor in a GLM analysis of activation measured across the whole brain.

#### Restricted Network Analysis

For our restricted network analysis, we selected a sub-set of ROIs based on prior knowledge of brain networks associated with adaptation (16, 20, 21). These ROIs included the bilateral primary motor cortices (M1), premotor cortices (PM), and primary sensory cortices (S1), the left thalamus, and the right cerebellar lobules 6 and 8 (CB6, CB8). The six bilateral cortical ROIs (L-M1, L-S1, L-PM, R-M1,R-S1, R-PM) formed a cortical sensorimotor network of interest, while the left motor ROIs, left thalamus, and right cerebellar ROIs formed a restricted corticothalamic-cerebellar network of the trained wrist (L-M1, L-PM, L-S1, L-Thalamus, R-CB6, R-CB8). Both networks were formed as a complete graph with 6 nodes (ROIs) and 15 edges (ROI pairs).

To test for the effect of condition (REST1, REST2, and REST3) on measured rsFC, we performed a mixed model analysis on each edge in our two networks of interest (sensorimotor and cortico-thalamic-cerebellar network). To test for changes associated with motor performance we compared model estimations of REST1 to REST2. To test for changes associated with dynamic motor control we compared REST1 to REST3. We additionally compared REST3 to the union of REST1 and REST2 to account for any effects due to the motor performance task or changes with time. To correct for multiple comparisons associated with the multiple edges within each network, we used a Bonferroni correction (*p*_*FWE*_ < 0.05), such that model results were deemed significant at *p <* 0.0033 (0.05/15 edges). Due to the conservative nature of this correction, edges that achieved a reduced model significance of *p*_*unc*_ < 0.01 are also reported, separately from effects with family wise error correction. At both significance levels, we corrected for the number of contrasts performed on model parameters, such that changes were considered significant at *p <* 0.0167 (0.05/3 contrasts).

#### Expanded Network Analysis

To supplement our restricted network analysis, we performed a network-based statistical analysis using all 65 task-related ROIs to identify any sub-networks of ROIs with significant change in resting state functional connectivity follow motor task performance (22). To test for network-level change in rsFC due to motor performance, we performed a paired t-test between REST1 and REST2. To test for changes due to dynamic motor control, we performed a paired t-test between REST1 and REST3. For all tests, initial connections between ROIs were thresholded at *t >* 3, and a topological false-discovery rate (*p*_*FDR*_ < 0.05) was applied to correct for the expected proportion of false discoveries among all clusters (sub-networks) over the entire network under the null hypothesis of no change.

#### ROI-to-Voxel Analysis

Finally, to supplement our ROI-ROI analyses, we performed a GLM analysis of three seed ROIs in the left sensorimotor network (L-M1, L-S1, L-PM) to resting state activity measured across the whole brain. In this analysis, the ROI’s average timeseries is regressed onto the timeseries of each voxel, to create subject level beta-maps of functional connectivity across the whole brain associated with the seed ROI. To quantify baseline rsFC associated with each seed ROI, we performed a group-level one-sample t-test on REST1 data. To identify clusters with significant change in rsFC between rest conditions associated with each seed ROI, we performed group-level paired t-tests between conditions. Comparison of REST1 and REST3 was used to test for effects of dynamic motor control and comparison of REST1 and REST2 was used to test for effects of motor execution. We chose a threshold of *p*_*unc*_ < 0.01 (at the voxel-level), *p*_*FDR*_ < 0.05 (at the cluster level) to test for significant effects. To correct for multiple comparisons (3 ROIs, 3 conditions) we used a voxel-wise threshold of *p*_*unc*_ < 0.0011 (*p <* 0.01/9, uncorrected) for each map.

#### Associations between rsFC and Behavior

To investigate associations between behavior on day one and rsFC measured in different resting state conditions, we performed a linear regression between behavior (MAE, MAI, RER) and rsFC measured within each restricted network of interest (sensorimotor, cortico-thalamic-cerebellar). We used the general model: *Y*_*behavior,D*1_ = *β*_0_ + *β*_1_*rsFC*_*REST* 1,*i*_ + *β*_2_*rsFC*_*REST* 2 −*REST* 1,*i*_ + *β*_3_*rsFC*_*REST*3_ − _*REST*1,*i*_ to identify edges *i* for further analysis that met one of the following criteria: a model significance of *p <* 0.05, any individual model parameter with a significance of *p <* 0.05. For edges that met these criteria, we performed a backwards, step-wise removal of non-significant model parameters until all the parameters in the model were significant (*p <* 0.05), or if the removal of further parameters resulted in a decrease in the model adjusted *R*^2^. To account for multiple comparisons associated with the multiple edges within each network, we used a Bonferroni correction (*p*_*FWE*_ < 0.05), such that model results were deemed significant if the reduced model reached a significance of *p <* 0.0033 (0.05/15 edges). Due to the conservative nature of this correction, edges that achieved a reduced model significance of *p*_*unc*_ < 0.01 are also reported, separately from those with family-wise error correction. At both significance levels, we considered model terms significant after correcting for the number of terms within the reduced model (*p*_*β,corr*_= 0.05/the number of terms in the reduced model). Associations between rsFC and behavior were then reported based on terms that reached significance within the reduced model.

#### Variance Decomposition

For edges with resting state functional connectivity that was significantly associated to behavior in our linear regression analysis, we performed an exploratory analysis using the MAD and AERS metrics to determine if observed effects in our rsFC conditions (REST1, REST2 − REST1, or REST3 − REST1) were driven by internal model formation processes (MAD), or alternate error reduction strategies (AERS). For day one associations, we used a linear model of the form: *Y*_*rsFC,i*_ = *β*_0_ + *β*_1_*M AD*_*D*1_ + *β*_2_*AERS*_*D*1_ to determine differential associations between rsFC (measured in a given edge *i* and resting state condition) and the two learning processes. For day two associations, we performed the same analysis to our day two variance decomposition metrics (i.e. *Y*_*rsFC,i*_ = *β*_0_ + *β*_1_*M AD*_*D*2_ + *β*_2_*AERS*_*D*2_). Because the MAD and AERS metrics are both normalized and independent from each other, significant model fits (p < 0.05) to nodes with larger *β*_1_ were determined to be associated with internal model formation, while nodes with larger *β*_2_ were determined to be associated with alternate error reduction strategies. For edges where the reduced model included functional connectivity in multiple conditions (e.g., *Y*_*behavior,D*1_ = *β*_0_ + *β*_1_*rsF C*_*REST* 1,*i*_ + *β*_2_*rsF C*_*REST* 2 *REST* 1,*i*_, |*β*_1_| > 0 and |*β*_2_| > 0 at *p <* 0.05), we investigated associations between each term (rsFC) and our control-specific metrics (MAD, AERS) independently. Post-hoc investigations were conducted to determine the direct associations between rsFC and learning strategy exhibiting the strongest effect (e.g., *Y*_*rsFC,i*_ = *β*_0_ + *β*_1_*M AD*_*D*1_) Additionally, for day two investigations, we performed a direct analysis of the reduced model form to MAD and AERS independently (e.g., *MAD*_*D*2_ = *β*_0_ + *β*_1_*rsF C*_*REST* 1,*i*_ + *β*_2_*rsF C*_*REST* 3 −*REST* 1,*i*_ + *β*_3_*M AD*_*D*1_), including the day one term to account for effects of previous behavior.

## ACKNOWLEDGMENTS

This work was supported by the AHA under award SDG no. 17SDG33690002, and by the NSF CBET under grant no. 1943712

